# Responses of the putative trachoma vector, *Musca sorbens*, to volatile semiochemicals from human faeces

**DOI:** 10.1101/741496

**Authors:** Ailie Robinson, Julie Bristow, Matthew V. Holl, Pateh Makalo, Wondu Alemayehu, Robin L. Bailey, David McLeod, Michael A. Birkett, John C. Caulfield, Virginia Sarah, John A. Pickett, Sarah Dewhirst, Vanessa Chen-Hussey, Christine M. Woodcock, Umberto D’Alessandro, Anna Last, Matthew J. Burton, Steve W. Lindsay, James G. Logan

**Affiliations:** Department of Disease Control, London School of Hygiene and Tropical Medicine, Keppel Street, London, WC1E 7HT, UK; Biological Chemistry and Crop Protection, Rothamsted Research, Harpenden, Hertfordshire, AL5 2JQ, UK; Medical Research Council Unit, The Gambia, The Gambia; The Fred Hollows Foundation, P.O. Box 6307, Addis Ababa, Ethiopia; Department of Clinical Research, London School of Hygiene and Tropical Medicine, Keppel Street, London, WC1E 7HT, UK; Department of Infectious Disease Epidemiology, London School of Hygiene and Tropical Medicine, Keppel Street, London, WC1E 7HT, UK; Global Partnerships Executive, The Fred Hollows Foundation, 12-15 Crawford Mews, York Street, London W1H1LX; ARCTEC, Chariot Innovations Ltd, London School of Hygiene and Tropical Medicine, Keppel Street, London, WC1E 7HT, UK; Department of Biosciences, Durham University, Durham, County Durham DH1 3LE, UK

## Abstract

**Background:** The putative vector of trachoma, *Musca sorbens*, prefers to lay its eggs on human faeces on the ground. This study sought to determine whether *M. sorbens* females were attracted to volatile odours from human faeces in preference to odours from the faeces of other animals, and to determine whether specific volatile semiochemicals mediate selection of the faeces.

**Methodology/Principal findings:** Traps baited with the faeces of humans and local domestic animals were used to catch flies at two trachoma-endemic locations in The Gambia and one in Ethiopia. At all locations, traps baited with faeces caught more female *M. sorbens* than control traps baited with soil, and human faeces was the most successful bait compared with soil (mean rate ratios 44.40, 61.40, 10.50 [*P*<0.001]; 8.17 for child faeces [*P*=0.004]). Odours from human faeces and some domestic animals were sampled by air entrainment. Extracts of the volatiles from human faeces were tested by coupled gas chromatography-electroantennography with laboratory-reared female *M. sorbens*. Twelve compounds were electrophysiologically active and tentatively identified by coupled mass spectrometry-gas chromatography, these included cresol, indole, 2-methylpropanoic acid, butanoic acid, pentanoic acid and hexanoic acid.

**Conclusions/Significance:** It is possible that some of these volatiles govern the strong attraction of *M. sorbens* flies to human faeces. If so, a synthetic blend of these chemicals, at the correct ratios, may prove to be a highly attractive lure. This could be used in odour-baited traps for monitoring or control of this species in trachoma-endemic regions.

**Author summary:** *Musca sorbens*, also known as the Bazaar Fly, visits people’s faces to feed on ocular and nasal discharge. While feeding, *M. sorbens* can transmit *Chlamydia trachomatis*, the bacterium that causes the infectious eye disease trachoma. Around 1.9 million people worldwide are visually impaired or blind from this disease. Although it is believed that *M. sorbens* transmits trachoma, very few studies have looked at ways to control this fly. A large-scale trial has shown that control of fly populations with insecticide reduces active trachoma disease prevalence. Odour-baited traps for the suppression of disease vector populations are an attractive option as there is no widespread spraying of insecticide, however, highly attractive baits are critical to their success. Here we demonstrate that the preference of these flies for breeding in human faeces is probably mediated by odour cues, and we isolate chemicals in the odour of human faeces that cause a response in the antennae of *M. sorbens*. These compounds may play a role in the specific attractiveness of human faeces to these flies, perhaps by being present in greater amounts or at favourable ratios. These may be developed into a chemical lure for odour-baited trapping to suppress *M. sorbens* populations.

## Introduction

The Bazaar Fly, *Musca sorbens*, is the putative vector of the blinding eye disease trachoma [1]. Adult *M. sorbens* feed on ocular and nasal secretions to obtain nutrition and liquid [2], and in doing so can transmit *Chlamydia trachomatis*, the bacterium that causes trachoma, from person to person. *Chlamydia trachomatis* DNA has been found on wild caught *M. sorbens* [3–5], and a laboratory study demonstrated mechanical transmission of *C. psittaci* between the eyes of Guinea Pigs by the closely related *Musca domestica* [6]. Strong evidence for the role of *M. sorbens* as vectors of trachoma comes from a cluster-randomised controlled trial that examined the impact of fly control interventions on trachoma prevalence [7]. Insecticide spraying significantly reduced the number of *M. sorbens* flies caught from children’s faces by 88 %, and a 56 % reduction in trachoma prevalence in children was observed. The provision of pit latrines, which by removing sources of open defecation controls *M. sorbens* juvenile stages, resulted in a 30 % decrease in flies on faces and a 30 % reduction in trachoma prevalence (non-significant).

These findings demonstrate that controlling the population density of *M. sorbens* may contribute to a decline in trachoma by reducing the number of fly-eye contacts, highlighting the disease-control potential of effective fly control tools. Odour-baited traps are receiving increased attention with regards to disease vectors, as the knowledge base around insect olfaction and attractive volatile chemicals expands [8–12]. Recent studies demonstrating the epidemiological significance of implementing chemical-based lures for vector-borne disease control [13] bolster the more widely accepted, and longstanding, tsetse fly example [14].

Mass deployment of an odour-baited trap for *M. sorbens*, based on the attractive volatiles in faeces, may suppress populations sufficiently to decrease the prevalence of trachoma. Alternatively, such traps could be used for entomological surveillance, and for monitoring and evaluating *M. sorbens* control programmes.

Female *M. sorbens* deposit their eggs on faeces, in which the larvae develop [15]. Previous studies have shown that *M. sorbens* preferentially breed in human faeces [2,16], and that flies emerging from human faeces are on average larger than those emerging from any other types of faeces, suggesting that human faeces may be an optimal larval development medium [16]. It is unknown, however, whether more prolific emergence from human faeces is due to better larval survival within human faeces, or due to more oviposition in human faeces caused by its relatively greater attraction to female flies. It is common for insects to use semiochemicals, volatile airborne chemical signals, to locate resources such as oviposition sites and to discriminate between resources of varying quality. It is therefore plausible that female *M. sorbens* visit human faeces more frequently relative to the faeces of other animals because of favourable semiochemical cues.

The aim of this study was to investigate the attractiveness of faeces from human beings and local domesticated animals to the Bazaar fly, *M. sorbens*. We conducted studies at two locations in The Gambia and one in Ethiopia, and sought to identify putative attractants in faecal odours.

## Methods

### Study areas and study periods

The study was carried out in The Gambia and Ethiopia. In The Gambia, there were two sites, the village of Boiram, Fulado West, Central River Division, The Gambia and the rural town of Farafenni. Studies were conducted in Boiram during the June-August 2009 rainy season, and in Farafenni in November/December 2009, immediately after the rainy season. The third site was Bofa Kebele (Oromia, central Ethiopia), identified as having highly prevalent active trachoma by the Global Trachoma Mapping Project. Here the study was conducted in February 2017 [17].

### Ethics

The Gambian study was approved by the Joint Gambian Government/Medical Research Council Laboratories Joint Ethics Committee (protocol number L2010.90, re: L2009.67, 01/12/09), and the Ethiopian study by the LSHTM ethics committee (reference number 11979/RR/5821) and the Oromia Regional Health Bureau Ethics Committee. In the Gambian study, written informed consent was provided by the heads of all compounds in which traps were sited or from which faeces were collected. In the Ethiopian study, written informed consent was provided by all participants including guardians of children.

### Trapping

#### Trap design

Fly traps were placed on the ground and consisted of a white plastic pot (7.8 cm high, 8.5 cm bottom diameter (D), 11.5 cm top D, Vegware, Edinburgh, UK) containing 50 g of faeces, soil, or left empty (Fig 1). Pots were covered by a lid (Gambian study, a disc cut from yellow sticky trap [Agrisense BCS Ltd, Pontypridd, UK]; Ethiopian study, the commercially available pot lid [Vegware] with yellow sticky trap stuck on [Agralan, Wiltshire, UK]) with a hole in the centre (Gambian study, 3.2 cm D, Ethiopian study, 4 cm D), covered on the underside with nylon mesh (Gambian study, 0.4 mm gauge [Lockertex, Warrington]; Ethiopian study, white polyester mesh, The Textile House). Traps were replaced daily.

**Fig 1.**
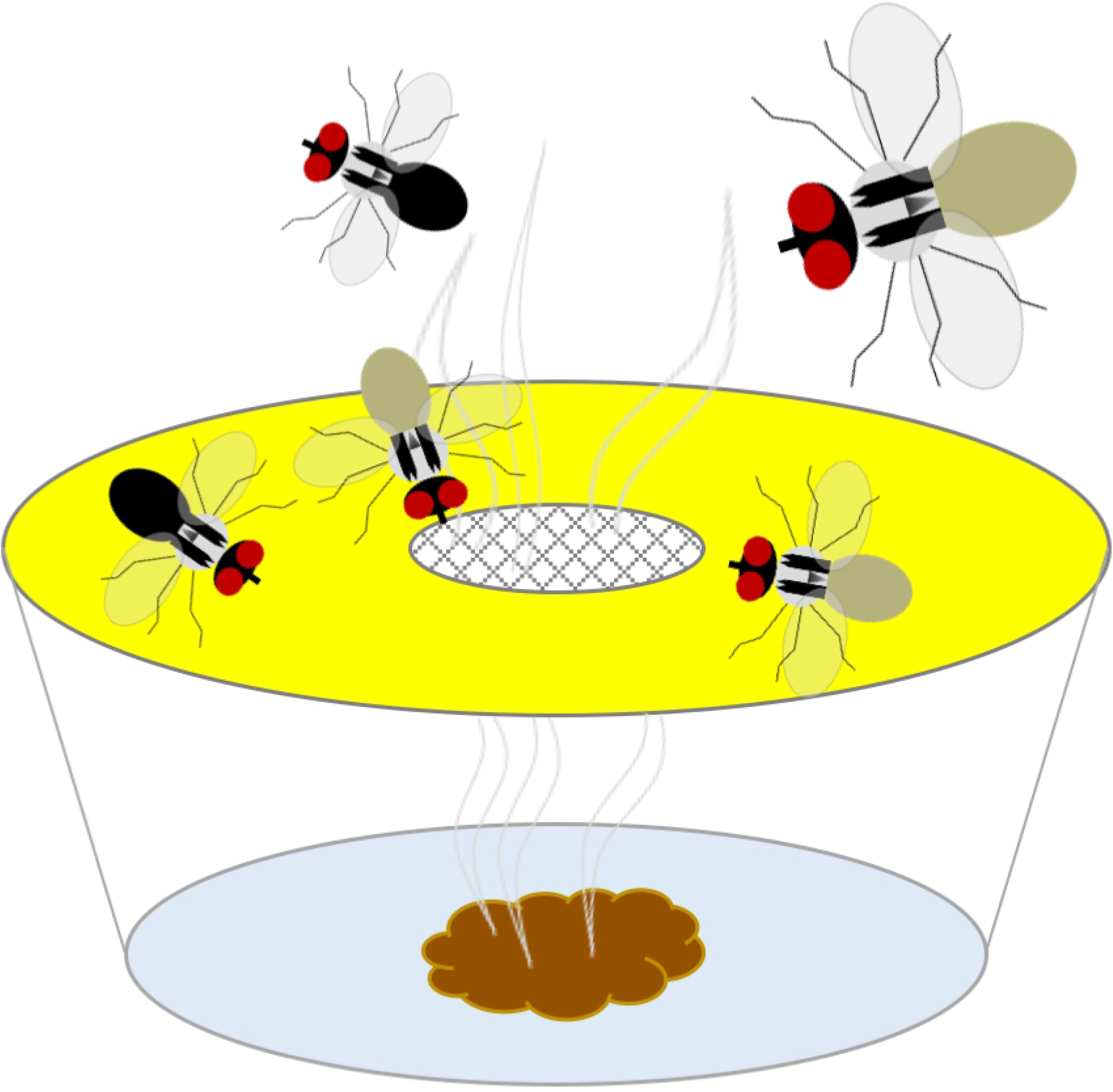
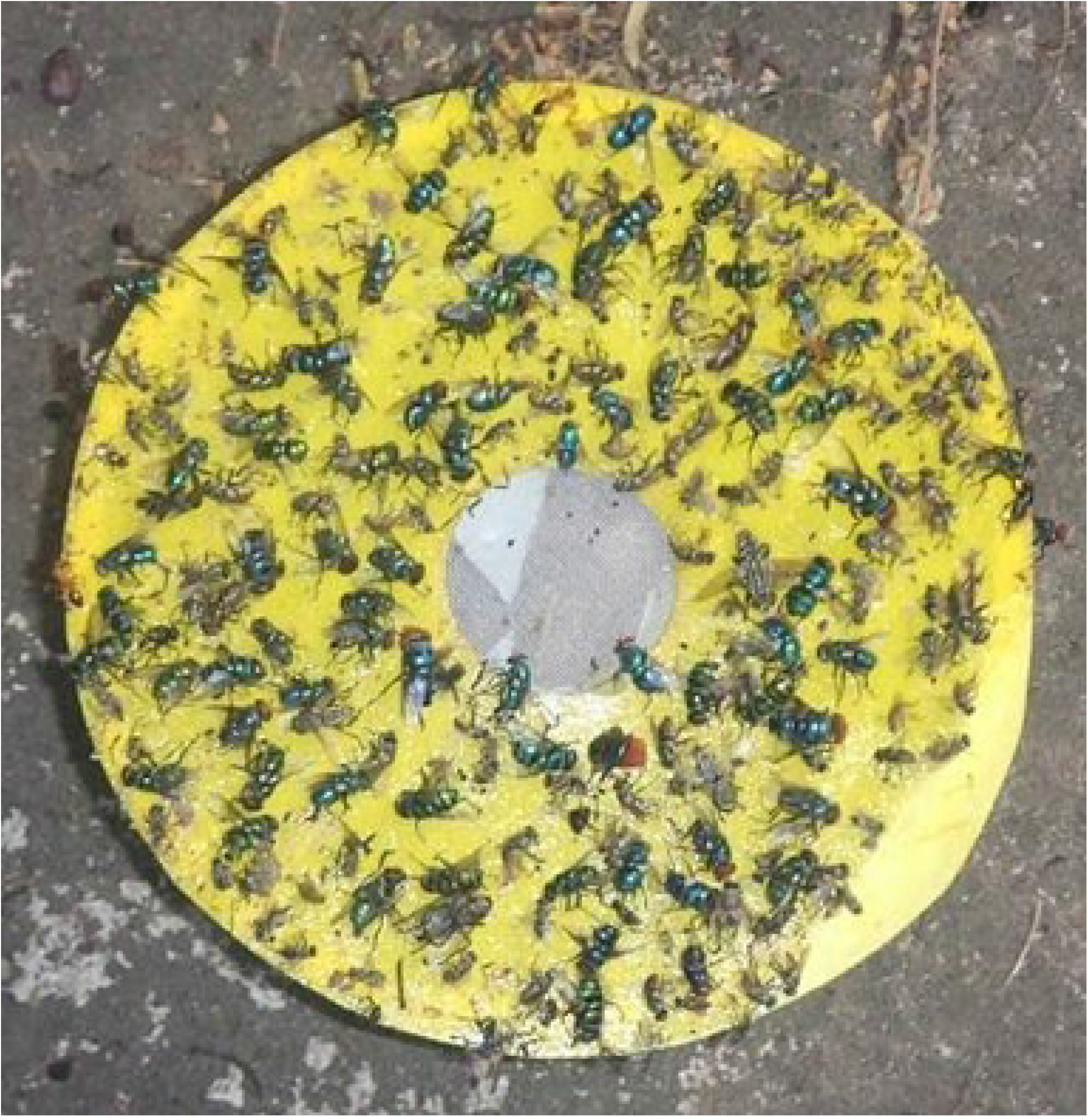
Bristow odour-baited trap for *Musca sorbens*. (A) Trap design, (B) typical fly catch on the yellow sticky discs on the trap.

In The Gambian study, a layer of glue (“Rat Stop”, 92 % polybutene, 8 % hexane) was added to the top of the yellow disc to increase trap catch. A thin band of “Rat Stop” was also applied to the side of each pot to prevent ants from gaining access to, and consuming, the trap catches. Wire frames, 38 cm^3^, coated in black electrical tape and with the upper side covered by blue plastic sheeting, were placed over the pot traps to protect against rain; in the Ethiopian study neither this protection nor the “Rat Stop” glue was used. A barrier of thorny acacia branches was placed in front of all traps to stop large animals interfering with them.

#### Collection of faeces bait

In the Gambian study, faeces (50 g, calf, cow, dog, donkey, horse, human and sheep) were collected for trap bait from open defecation in the compounds between 07:00 h and 11:30 h and weighed on a balance (Salter ARC 1066, accurate to ± 1 g). In Boiram, human faeces were obtained from two adjacent compounds, the children of which defaecated on the ground. In Farafenni, human faeces were obtained from a compound with five children between the ages of eight and 15 years, who defaecated in a plastic potty in the morning. Cow and calf faeces were collected from areas where cattle were confined at night. Cattle grazed in the bush, calves were under one-year-old and fed on milk. Dog faeces were unavailable in Boiram. Donkeys and horses were fed cous (millet), supplemented in the village by grazing. Their faeces were collected from family compounds. Sheep faeces were collected from shelters where the animals were kept overnight. Sheep in the village grazed in the bush, whilst those in town scavenged in the streets. Tobaski, a religious festival during which rams or sheep are slaughtered, occurred between the first and second experiments. Before Tobaski, sheep had their diets supplemented with milk, millet porridge or Senegalese feed blocks known as *repas* of unknown composition. After Tobaski the sheep had a less nutritious diet. Soil samples were taken from uncontaminated areas close to the trap site and used as control bait. In the Ethiopian study, human adult and human child, cow and donkey faeces (50 g) were collected from the compounds where open defecation is commonplace, and weighed (Ascher Portable Digital Scale, accurate to 0.01 g).

#### Experimental design and data analysis

Traps containing faeces or soil bait were set daily, 50 cm apart, along a transect. Latin Square (LS) designs were used so that baits were rotated between trap positions daily, allowing trapsite variation and bais, and preventing baits having an effect on adjacent traps. In the Gambian study, eight traps were set daily (seven faeces baits and soil, with an empty pot instead of dog faeces in Boiram) and the eight by eight LS was repeated twice in each site (*n*=16 trap days per location). In the Ethiopian study, a transect of five traps (four faeces and one soil bait) was set on day one outside one household, and thereafter five-trap transects were deployed outside two (new) households daily until day six, giving a total of 11 trapping days per treatment (trap/bait). The position of the traps within each transect was rotated according to a LS design. Environmental variables were recorded as follows: Bioram, max/min ambient temperature, presence/absence of precipitation (thermometers hung nearby and daily observation of precipitation); Farafenni max/min ambient temperature and humidity (TinytagPlus datalogger, Gemini dataloggers); Oromia, start/finish ambient temperature, start/finish ambient humidity (Colemeter thermometer hygrometer humidity meter).

After 24 h, *M. sorbens* found adhering to the sticky trap lids were identified according to taxonomic keys [18,19] and counted. In The Gambian study, *M. sorbens* were counted, females dissected for gravidity and *Musca domestica* were counted. In the Ethiopian study, *M. sorbens* were counted and sexed. Negative binomial regression, which accounted for the over-dispersed nature of trap catch data, was used to model the relationship between trap bait and the number of flies caught. The effect of trap bait on the likelihood that a trapped *M. sorbens* was female was analysed using logistic regression (analyses performed in Stata [v. 15, StatCorp]).

### Chemistry of bait attraction

#### Collection of volatiles

Air entrainment of faeces samples was performed only during the Gambian study. Five human faeces samples were collected (50 g) from Farafenni and Boiram into individual sterile polyethyleneterephthalate (PET) cooking bags (Sainsbury’s Ltd, UK). Volatiles from the faeces were collected using a portable air entrainment kit. The bag containing the sample was sealed to an aluminium disc with air inlet and outlet holes using bulldog clips. The air entrainment kit, comprising an inflow and outflow pump and charcoal filters (VWR Chemicals BDH, 10-14 mesh, 50 g, preconditioned under a stream of nitrogen at 150 °C for a minimum of two hours), was connected to the bag and air inflow (16 L/min) set higher than outflow (2 x 7 L/min), creating positive pressure to prevent entrance of environmental volatiles. PTFE tubing and rubber ferrules were used for all connections. The apparatus was cleaned before and after use with ethanol (100 %, Sigma-Aldrich, Gillingham, UK). Volatiles were collected in the outflow onto Porapak Q polymer (50 mg, mesh size 50/80, Supelco), contained inside a glass tube and held with two plugs of sterile silicanised glass wool (‘Porapak tubes’). These had been conditioned prior to use by repeated washing with redistilled diethyl ether and heating to 132 °C for 2 h under a stream of constant (filtered) nitrogen. After 12 h of volatile collection, Porapak tubes were sealed in ampoules under filtered nitrogen for transport and storage. Ampoules were initially stored at the study site at 4 °C (for a maximum of six weeks), then in the UK volatiles were eluted from the Porapak tubes using freshly-distilled diethyl ether (‘extract’) and stored in vials at −20 °C until analysis.

#### Gas chromatography

Air-entrainment extracts were concentrated (from 750 to 50 μL), and samples (1 μL) analysed by gas chromatography (GC). The GC (Agilent 6890N, equipped with a temperature programmable on-column injector, flame ionization detector [FID] and using hydrogen as a carrier gas) was fitted with a non-polar HP-1 column (10 m, 0.53 mm internal diameter (ID), 2.65μm film thickness). The following GC program was used: oven temperature maintained at 30 °C for 2 min, increased by 10 °C per min to 230 °C, then held at 230 °C for 30 minutes.

#### Fly rearing

After counting and identification, live female *M. sorbens* were collected from the sticky traps with blunt forceps and placed into Insect Rearing Cages (BugDorm, 47.5 cm^3^) containing water (soaked blue roll), white sugar cubes and either human faeces (20 g on a small pot of damp soil) or full fat Ultra-high temperature processing (UHT) milk (soaked into cotton wool). The latter served as an oviposition medium and protein source respectively [20], and every third day larvae were transferred onto larval diet medium: molasses sugar (8 g), dried yeast (7 g), full fat UHT milk (100 mL), water (200 mL) and wheat bran. Cages containing artificial diet were kept indoors in The Gambia at 25-28 ° C and 25-50 % relative humidity (RH), while cages containing faeces were kept in natural daylight in a ventilated outbuilding, at 17-44 ° C and 8-65 % RH, with a roughly 12:12 light/dark photo-period. Six days after they were placed in the cages, two samples of artificial larval diet and one sample of faeces were removed from oviposition cages and the soil carefully scraped away to expose the pupariae. Emerging adults were used for GC-EAG.

#### Coupled gas chromatography-electroantennography (GC-EAG)

Randomly selected *M. sorbens* females from the laboratory-reared colony were chilled on ice until movement ceased, the heads severed, and abdomens dissected to determine gravidity. One antenna was removed to reduce noise in the recording. The tip of the other antenna was cut off before inserting the antenna into a glass electrode containing a silver/silver chloride wire and filled with Ringer’s solution (7.55 g NaCl, 0.64 g KCl, 0.22 g CaCL_2_, 1.73 g MgCl_2_, 0.86 g Na_2_HCO_3_, 0.61 g Na_3_PO_4_/L distilled water). Another electrode was inserted into the back of the flies’ head to complete the circuit. This assemblage was held in a constant stream (1 L/min) of humidified, charcoal-filtered air. A sample of the extract (1 μL of a representative 50 μL concentrated 12-h air entrainment extract) was injected onto a 30 m non-polar polydimethylsiloxane (HP1) column (internal diameter 0.32 mm, solid phase thickness 0.52 μm) in a gas chromatograph (Hewlett Packard HP6890 with a cool-on-column injector, hydrogen carrier gas and flame ionisation detector). The following GC program was used: oven temperature maintained at 40 °C for 2 min, increased by 5 °C per min to 100 °C, then raised by 10 °C per min to 250 °C. Emerging compounds were delivered simultaneously to the flame ionisation detector and the airstream blowing over the antenna. The signal was amplified 10,000 times by the Intelligent Data Acquisition Controller-4, and signals were analysed by using EAD 2000 software (both Syntech, The Netherlands). Antennal responses were correlated visually to compound peaks by overlaying traces on a light box, and the procedure repeated for four female flies.

#### Coupled gas chromatography-mass spectrometry (GC-MS)

Compounds found to be electrophysiologically-active were identified by GC-MS only. One concentrated extract was diluted tenfold prior to injecting (1 μL) onto a HP1 column (dimensions as for GC-EAG; Hewlett HP 5890 GC fitted with a cool-on-column injector, helium carrier gas and FID, with a deactivated HP1 pre-column [0.53 mm ID]). The following program was used: oven temperature maintained at 30 °C for 5 min, increased by 5 °C per min to 250 °C. A VG Autospec double-focusing magnetic sector mass spectrometer (MS) using electron impact ionisation (70 eV, 250 °C) was coupled to the GC and data analysed using an integrated data system (Fisons Instruments, Manchester, UK). Compounds were identified using the NIST 2005 database of standards (NIST/EPA/NIH mass spectral library version 2.0, Office of the Standard Reference Data Base, National Institute of Standards and Technology, Gaithersburg, Maryland).

## Results

### Trapping

A total of 1734 muscid flies were caught in Boiram across all traps between 17^th^ July 2009 and 7^th^ August 2009. Of these, 382 were *M. sorbens*, (82.9 % female), 1046 M. *domestica* (69.7 % female) and 306 unidentified *Musca* spp. (80.3 % female). A total of 1899 flies were caught in Farafenni between 18^th^ November 2009 and 10^th^ December 2009, of which 1754 were *M. sorbens*, (96.5 % female) and 145 *M. domestica* (86.2 % female). Aside from horse or sheep faeces, for all other faeces baits more than 60 % of flies collected were gravid. A total of 152 *M. sorbens* were caught between 19^th^ and 27^th^ February 2017 in Oromia, of those that could be sexed (96.1 %), most were female (90.4 %). Gravidity was not measured, and other flies/arthropods not counted. The distribution of *M. sorbens* trap catches per night by bait type are shown in Fig 2, S1-S2 Tables.

**Fig 2.**
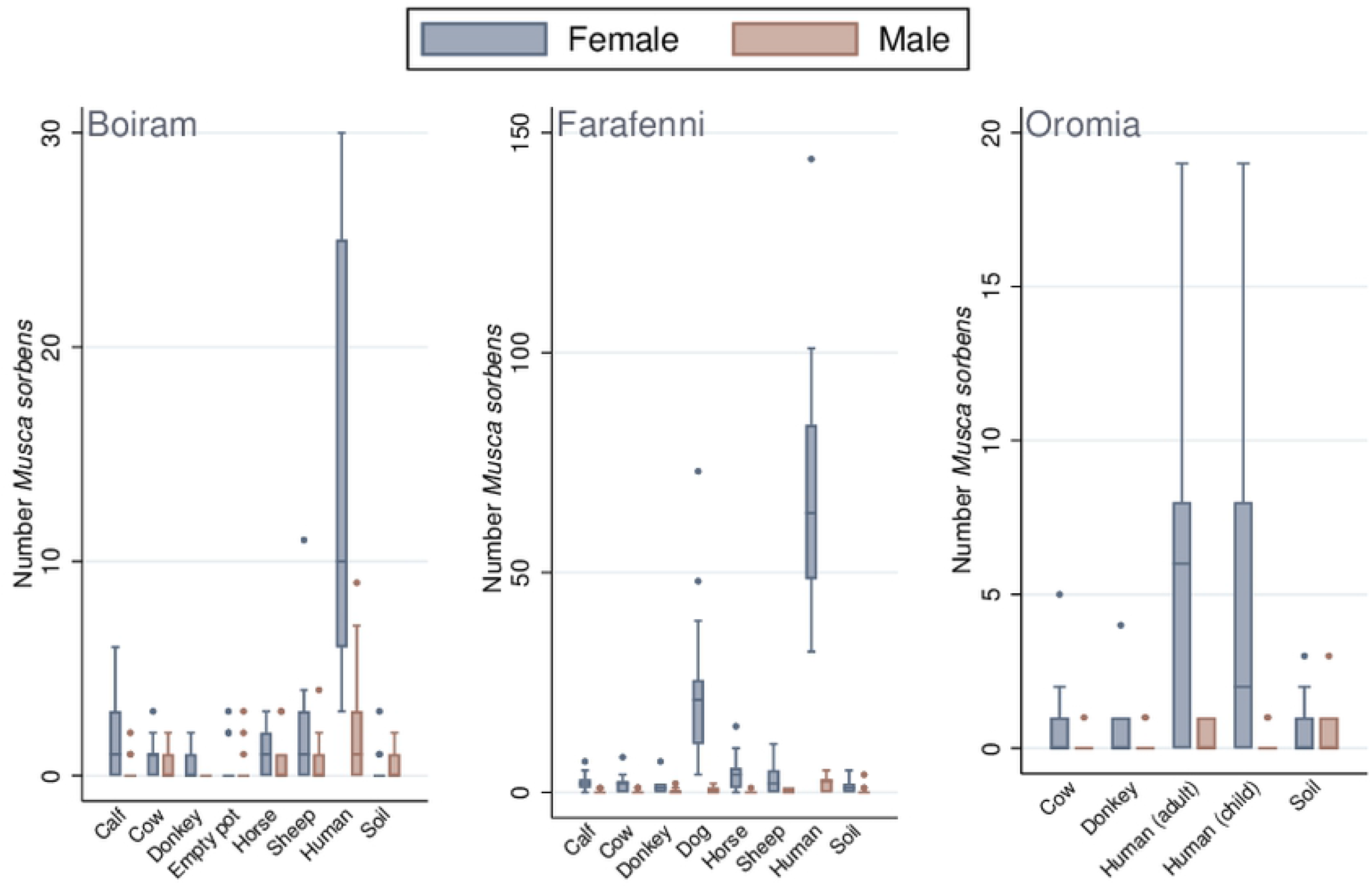
Male and female *Musca sorbens* caught by different faeces bait, in the three studies. (boxes, median and interquartile range; points, outliers).

In all three studies, there was an association between the type of trap bait and the number of female *M. sorbens* caught (*P*<0.001, Table 1). Human faeces were found to be the most attractive bait to female *M. sorbens*, although, when using soil bait as the baseline, the estimated rate ratios varied substantially between studies. In Boiram the mean rate ratio (RR) was 44.4 (95 % confidence intervals [CI] 15.5-127.3, *P*<0.001), in Farafenni 61.4 (95 % CI 32.3-117.0, *P*<0.001) and in Oromia 10.5 (95 % CI 2.5-43.5, *P*=0.001) in traps baited with adult faeces and 8.2 (95 % CI 2.0-34.0, *P*=0.004) when child faeces were used (Table 1).

**Table 1.** *Musca sorbens* caught by different baits in three study sites

|  |  | Female *Musca sorbens* |  |  | Male *Musca sorbens* |  |  |
| --- | --- | --- | --- | --- | --- | --- | --- |
| **BOIRAM** |  | Rate ratio (95 % CI) | P-value<sup>A</sup> | P-value<sup>B</sup> | Rate ratio (95 % CI) | P-value<sup>A</sup> | P-value<sup>B</sup> |
| Bait | Calf faeces | 4.60 (1.50-14.12) | 0.008 | <0.001 | 0.80 (0.61-4.13) | 0.790 | 0.086 |
|  | Cow faeces | 2.20 (0.66-7.31) | 0.198 |  | 1.00 (0.21-4.86) | 1.000 |  |
|  | Donkey faeces | 1.40 (0.39-5.04) | 0.607 |  | <sup>C</sup> |  |  |
|  | Empty pot | 1.40 (0.39-5.04) | 0.607 |  | 1.20 (0.26-5.60) | 0.817 |  |
|  | Horse faeces | 3.20 (1.01-10.15) | 0.048 |  | 1.80 (0.41-7.82) | 0.433 |  |
|  | Human faeces | 44.40 (15.49-127.30) | <0.001 |  | 5.60 (1.43-21.96) | 0.014 |  |
|  | Sheep faeces | 5.20 (1.71-15.83) | 0.004 |  | 1.60 (0.36-7.08) | 0.536 |  |
|  | Soil | 1.0 |  |  | 1.0 |  |  |
| Climate | Av. temp<sup>D,E</sup>, continuous | 1.15 (0.92-1.45) |  | 0.225 | 1.45 (1.08-1.94) |  | 0.014 |
|  | Rainfall<sup>F,E</sup>, Yes | 1.21 (0.40-3.62) |  | 0.737 | 0.82 (0.21-3.21) |  | 0.779 |
| **FARAFENNI** |  |  |  |  |  |  |  |
| Bait | Calf faeces | 2.00 (0.97-4.11) | 0.059 | <0.001 | 0.33 (0.06-1.78) | 0.198 | <0.001 |
|  | Cow faeces | 1.70 (0.83-3.58) | 0.145 |  | 0.50 (0.12-2.17) | 0.355 |  |
|  | Dog faeces | 20.70 (10.80-39.57) | <0.001 |  | 1.17 (0.35-3.85) | 0.800 |  |
|  | Donkey faeces | 1.20 (0.57-2.63) | 0.607 |  | 0.83 (0.23-3.00) | 0.781 |  |
|  | Horse faeces | 3.70 (1.85-7.28) | <0.001 |  | 1.17 (0.02-1.46) | 0.106 |  |
|  | Human faeces | 61.40 (32.26-117.02) | <0.001 |  | 5.33 (1.97-14.47) | 0.001 |  |
|  | Sheep faeces | 2.50 (1.22-5.08) | 0.012 |  | 0.89 (0.25-3.21) | 0.857 |  |
|  | Soil | 1.0 |  |  | 1.0 |  |  |
| Climate | Av. temp<sup>G</sup>, continuous | 1.11 (0.96-1.27) |  | 0.150 | 1.26 (1.05-1.52) |  | 0.015 |
|  | Av. RH<sup>F</sup>, Yes | 1.02 (0.98-1.06) |  | 0.294 | 1.02 (0.97-1.07) |  | 0.412 |
| **OROMIA** |  |  |  |  |  |  |  |
| Bait | Cow faeces | 1.33 (0.20-6.35) | 0.718 | <0.001 | 0.17 (0.02-1.42) | 0.102 | 0.359 |
|  | Donkey faeces | 1.00 (0.21-5.01) | 1.000 |  | 0.33 (0.06-1.71) | 0.189 |  |
|  | Adult human faeces | 10.50 (2.54-43.48) | 0.001 |  | 0.50 (0.12-2.09) | 0.342 |  |
|  | Child faeces | 8.17 (1.96-34.03) | 0.004 |  | 0.33 (0.06-1.71) | 0.189 |  |
|  | Soil | 1.0 |  |  | 1.0 |  |  |
| Climate | Av. temp<sup>G</sup>, continuous | 0.89 (0.70-1.14) |  | 0.366 | 1.48 (1.08-2.04) |  | 0.015 |
|  | Av. RH<sup>F</sup>, Yes | 1.01 (0.96-1.07) |  | 0.721 | 1.09 (1.02-1.16) |  | 0.007 |

<sup>A</sup> P-value comparing this category with baseline (soil)
<sup>B</sup> P-value testing hypothesis that bait is associated with number of flies trapped
<sup>C</sup> Value could not be estimated as no male *Musca sorbens* caught in traps baited with donkey faeces
<sup>D</sup> Adjusted for rainfall
<sup>E</sup> Based on 13 days data (two days missing)
<sup>F</sup> Adjusted for average temperature
<sup>G</sup> Adjusted for average RH

For non-human faeces baits, in Farafenni, dog, horse and sheep faeces caught more female *M. sorbens* than soil-baited traps (dog, RR 20.7, 95 % CI 10.8-39.6, *P*<0.001; horse, RR 3.7, 95 % CI 1.9-7.3, *P*<0.001; sheep, RR 2.5, 95 % CI 1.2-5.1, *P*=0.012 Table 1). In Boiram, traps baited with horse, sheep and calf faeces were more attractive than soil-baited traps (RR 3.2, 95 % CI 1.0-10.2, *P*=0.048; RR 5.2, 95 % CI 1.7-15.8, *P*=0.004; RR 4.6, 95 % CI 1.5-14.1, *P*=0.008 respectively, Table 1). Dog faeces were not tested in Boiram or Oromia. In Oromia no evidence was found of a difference in rates between the non-human faeces baits (cow or donkey) and the soil bait.

Further to being more attractive than the soil bait, human faeces were found to be more attractive than the second most attractive bait at each site (*P*<0.001). In Boiram, human faeces-baited traps caught 8.5 times as many female *M. sorbens* (95 % CI 4.2-17.2) than sheep faeces-baited traps. In Farafenni, the rate ratio was 3.0 (95 % CI 1.9-4.7) compared to dog faeces-baited traps, and in Oromia, RR 7.9 (2.0-30.8) and RR 6.1 (1.6-24.1) in human and child faeces-baited traps respectively, compared to cow faeces-baited traps.

In all studies, there were increased odds that *M. sorbens* caught by human faeces bait relative to those caught in the soil control would be female (Boiram, odds ratio [OR] 7.93 [95 % CI 2.16-29.1) *P*=0.002; Farafenni, OR 11.52 [95 % CI 4.29-30.96) *P*<0.001; Oromia adult, OR 14 [95 % CI 3.63-53.99) *P*<0.001; Oromia child, OR 32.67 [95 % CI 5.59-191.06) *P*<0.001; S3 Table). Similarly, increased odds for female *M. sorbens* being caught were observed for dog and horse faeces baits in Farafenni, and cow faeces bait in Oromia (S3 Table).

### Coupled gas chromatography-electroantennography

Twelve compounds from the headspace of the human faecal sample elicited an antennal response from two or more *M. sorbens* females. The compounds were subsequently identified by GC-MS as 3-ethylpentane, 2-methylpropanoic acid, butanoic acid, pentanoic acid, hexanoic acid, cresol, 2-phenylethanol, valerolactam, dimethyl tetrasulphide, indole, 2-dodecanone and an unidentified cholesterol derivative (Table 2).

**Table 2.** Tentative identification of compounds eliciting electrophysiological responses in *Musca sorbens* females

| Peak number | Compound identified |
| --- | --- |
| 1 | 3-Ethylpentane |
| 2 | 2-Methylpropanoic acid |
| 3 | Butanoic acid |
| 4 | Pentanoic acid |
| 5 | Hexanoic acid |
| 6 | Isomer of cresol |
| 7 | 2-Phenylethanol |
| 8 | Valerolactam |
| 9 | Dimethyl tetrasulphide |
| 10 | Indole |
| 11 | 2-Dodecanone |
| 12 | Cholesterol derivative |

## Discussion

We found evidence that *M. sorbens*, the putative vector of trachoma, is strongly attracted to odours produced by human faeces, which attracted and caught the greatest number of *M. sorbens* in all three studies. This was particularly so with female *M. sorbens* relative to males. In the Gambian study, the majority of females were found to be gravid. These findings confirm that female *M. sorbens* are attracted to faeces for oviposition, as is widely accepted [2,15,21], and further demonstrate that volatile cues alone are responsible for this attraction as the faeces baits could only have been visible to flies if they were very close, directly above, and could see down through the mesh. A previous study conducted in The Gambia [16] demonstrated attraction to human faeces, but because the faeces were not hidden from the flies, visual cues could not be ruled out as a stimulus. In the current study, there was also evidence that some of the non-human faeces baits were relatively more attractive than others (e.g. calf and sheep in Boiram attracted more female *M. sorbens* than the soil control, as did dog and horse in Farafenni). The high rate ratio for female *M. sorbens* caught using dog faeces in Farafenni (the only site with dogs) may indicate a preference for non-herbivore faeces. The mean rate ratio of *M. sorbens* females caught using human faeces relative to the soil control varied between the three sites, but in all cases a large effect was seen. The Gambian studies were conducted in two different ecological settings and at different times of the year, making it impossible to account for the differences in fly abundance between sites. At the Oromia site, trap catch may have been removed by birds or other insectivores as no protective wire frames were used. Even more so, the environment, including local abiotic and biotic factors, and *M. sorbens* population in Ethiopia is likely to be so different to that in The Gambia that comparison of population density between studies and based on a small sampling window would not be appropriate.

Differential attraction to different types of faeces could be due to the presence or absence of certain semiochemicals that attract or repel the flies. Of the *M. sorbens* EAG-active compounds tentatively identified here, short chain fatty acids (SCFA, including 2-methyl propanoic acid, butanoic acid, pentanoic acid and hexanoic acid), indoles, cresols and sulphur compounds are known volatile organic compounds in faeces [22–25]. SCFA are produced in the gut by bacterial fermentation of carbohydrates and proteins [26], and are commonly found in vertebrate associated volatiles including urine and faeces [22,27–30]. Their detection by host-seeking arthropods is well documented [31–35]. The aromatic compounds identified (cresol, 2-phenylethanol, indole) are likely to be fermentation products of the aromatic amino acids tyrosine, phenylalanine and tryptophan [24].

Several of these faeces-associated volatiles have been described in similar entomological studies. Antennal response by *M. domestica* (GC-EAG) detected nine compounds in pig faeces volatiles, with butanoic acid and indole eliciting strong responses in subsequent dose-response EAG [36]. Both *m*- and *p*-cresol in the headspace of canine faeces elicited antennal response in the common green bottle fly, *Lucilia sericata*, and a chemical blend including these compounds was as attractive to flies as the faeces [37]. Cresols (isomer not identified) in the volatiles from rat carrion also elicited antennal response from *L. sericata* [38], and female and male *M. domestica* responded (by EAG) to volatiles including butanoic acid, hexanoic acid, 2-phenylethanol and *p*-cresol in the headspace of vinegar [39]. Similarly, butanoic acid and *p*-cresol were among chemostimulants of *Stomoxys calcitrans* detected in the headspace of rumen volatiles, these were also found to elicit activation and attraction in a wind tunnel [40]. Microbial degradation is thought to lead to the production of *m*- and *p*-cresol in cattle urine, again found to elicit an EAG response in *S. calcitrans* [41]. *Stomxys calcitrans* is a muscid fly with coprophagous larvae, known to be attracted to faecal odours. This fly has been shown to select faeces by their odour [42], as demonstrated here with *M. sorbens*, and the chemostimulant compounds thought to be responsible for that attraction included butanoic acid, indoles, *p*-cresol and sulphides.

Taken together, the faecal semiochemicals described here are commonly isolated and/or detected as they are products of bacterial decomposition, and as such are frequently detected by filth flies, most likely as cues for oviposition sites. To underpin the specific attractiveness of human faeces to *M. sorbens* therefore, variation in amounts emitted, or ratios of compounds present, must distinguish this oviposition medium.

## Conclusion

Our study demonstrates that female *M. sorbens* at three different study locations, in both West Africa and East Africa, are preferentially attracted to the volatiles of human faeces, as evidenced by attraction in the absence of visual cues. We provide evidence that twelve compounds are putative attractants that may play a role in this response, by identifying, for the first time, compounds including short chain fatty acids and aromatic compounds that are detected by the antennae of *M. sorbens*. Further work is required to optimise chemical blends and release rates, to produce a synthetic lure to which the behavioural responses of *M. sorbens* can be investigated. Establishing those with attractive properties may lead to the design of baits for odour-baited traps, which could be used for *M. sorbens* surveillance or even population suppression or control.

## Acknowledgements

The authors thank all of the families and communities at the study sites for allowing us to conduct this work around their homes. We are particularly grateful for the contributions of the MRC Gambia fieldworkers Pateh Makalo, Kemo Ceesay and Dembo Camara, without whose hard work, dedication and ingenuity this study would not have been possible, and for the assistance of Keol Debele, the entomology fieldworker in Oromia. We are also grateful for the assistance from Aida Abashawl in project co-ordination, Nazif Jemal in communications, and Munira Haji Mohammed Yusuf, Helen W/Semayat, and Demitu Legesse in translation in the Oromian work.

## Author contributions

**Conceptualization:** RB, JAP, SD, VCH, SWL, JGL

**Data Curation:** JB

**Formal Analysis:** AR, DM, JB

**Funding Acquisition:** SWL, JGL, MJB, AL, VS

**Investigation:** JB, PM, MVH, JCC, JAP, CW

**Methodology:** JB, SWL, JGL

**Project Administration:** JB, MVH, WA, VCH, SD, AL, VS Resources: WA, MAB, UDA, VS

**Supervision:** MAB, JCC, JAP, SD, VCH, UDA, SWL, JGL, AL, MJB

**Visualization:** JB, AR, DM

**Writing - Original Draft Preparation:** AR, JB, JGL

**Writing - Review & Editing:** AR, JB, MVH, PM, WA, RB, DM, MB, JCC, VS, JAP, SD, VCH, CW, UDA, MJB, AL, SWL, JGL

## Supporting Information

**S1 Table. Median female *Musca sorbens/trap/night* (IQR)**

**S2 Table. Median male *Musca sorbens/trap/night* (IQR)**

**S3 Table. Odds of *Musca sorbens* caught by different bait types being female**

**S4 dataset. *Musca sorbens* trapping dataset (raw)**

